# Parental environmental effects are common and strong, but unpredictable, in *Arabidopsis thaliana*

**DOI:** 10.1101/2021.11.04.467350

**Authors:** Vít Latzel, Markus Fischer, Maartje Groot, Ruben Gutzat, Christian Lampei, Joop Ouborg, Madalin Parepa, Karl Schmid, Philippine Vergeer, Yuanye Zhang, Oliver Bossdorf

## Abstract

The phenotypes of plants can be influenced by the environmental conditions experienced by their parents. In some cases, such parental effects have been found to be adaptive, which has led to much speculation about their ecological and evolutionary significance. However, there is still much uncertainty about how common and how predictable parental environmental effects really are. We carried out a comprehensive test for parental effects of different environmental stresses in the model plant *Arabidopsis thaliana*. We subjected plants of three *Arabidopsis* genotypes to a broad range of biotic or abiotic stresses, or combinations thereof, and compared their offspring phenotypes in a common environment. The majority of environmental stresses (16 out of 24 stress treatments) caused significant parental effects, in particular on plant biomass and reproduction, with positive or negative effects ranging from −35% to +38% changes in offspring fitness. The expression of parental effects was strongly genotype-dependent, with some effects only present in some genotypes but absent, or even in the opposite direction, in others. Parental effects of multiple environmental stresses were often non-additive, and their effects can thus not be predicted from what we know about the effects of individual stresses. Intriguingly, the direction and magnitude of parental effects were unrelated to the direct effects on the parents: some stresses did not affect the parents but caused substantial effects on offspring, while for others the situation was reversed. In summary, parental environmental effects are common and often strong in *A. thaliana*, but they are genotype-dependent and difficult to predict.

**Significance:** Stress experienced by plants can alter the phenotypes of their offspring. To understand the ecological and evolutionary significance of such parental effects, we must know how common and how predictable they are. In a large experiment with *Arabidopsis thaliana*, we show that the majority of 24 environmental stresses cause significant, and often strong, positive or negative parental effects. However, we also find that parental effects are genotype-specific and unrelated to the direct effect of individual stresses, and that multiple stresses often act in non-additive ways across generations. Thus, parental effects appear to be common and strong, but difficult to predict. Our findings have important implications for the study of plant responses to environmental change, and the design of stress experiments.

## Introduction

Phenotypic variation is at the heart of ecology and evolution. The variation in phenotype that we observe among individuals of the same species either reflects underlying genetic differences, and thus the evolutionary potential of a species, or it results from plastic responses to the environment, and could thus be related to a species’ environmental tolerance. A third source of phenotypic variation are parental effects, where the environmental conditions of parents affect the phenotypes of their progeny (1, 2, 3). Parental effects are somewhat peculiar in that they can generate patterns of resemblance among relatives that would usually be considered evidence for underlying genetic variation, while in fact they represent special cases of phenotypic plasticity that extend across generations. The biological mechanisms that cause parental effects include simple nutritional effects such as differential seed provisioning, but also physiological effects mediated by hormones, toxins or other cytosol components, or even epigenetic mechanisms where differential DNA methylation or chromatin changes are passed on to offspring (2, 4, 5).

Previous studies showed that parental effects can be ecologically important (e.g. 6, 7, 8) and also influence evolution (e.g. 9; 10, 11, 12, 13). In particular the demonstration that some parental effects are adaptive, with offspring thriving better in parental than non-parental environments (e.g. 6, 14, 15, 16, 17, 18), triggered a debate to what extent parental effects may be evolved mechanisms and a means of rapid adaptation to environmental change (e.g. 2, 12, 19, 20, 21, 22). However, despite great current interest in parental effects, many important questions remain unresolved.

One of the key challenges in the study of parental effects is to understand how general and how strong they really are. An increasing number of studies showed that parental effects can be substantial, and that they can both increase or decrease offspring fitness (e.g. 6, 8, 23, 24, 25, 26, 14, 18, 27, 28, 29), but many of these studies tested a single environmental factor on a single species, sometimes using only a single genotype (but see e.g. 17, 28, 30, 31, 32, 33). As a consequence, we still do not have a good idea of how widespread parental effects are across different environmental factors, and how consistent they are across species and genotypes. Given that non-successful tests for parental effects are more likely to end up in file drawers, researchers sceptical of parental effects might suspect that studies as the ones cited above merely represent ‘freak’ cases that cannot be generalized. Ultimately, the debate can only be settled through comprehensive experiments that test for parental effects across multiple species, genotypes and/or environmental factors.

Another fundamental question about parental effects is how predictable they are. For instance, is the magnitude and direction of a parental effect related to (and thus predictable from) the direct effect of an environmental stress on the parental generation? Intuitively, one should expect that environmental factors with stronger effects on parents are more likely to also affect their offspring, and that environmental factors with little or no effects on the parents should neither affect their offspring. But is this really true? We are not aware of any published study that has tested these simple but important assumptions.

Environmental change usually involves simultaneous changes in multiple environmental factors (34, 35, 36, 37). Still, most previous studies on parental effects worked with single environmental factors. We know, however, that the direct effects of multifactorial environmental changes are often non-additive (e.g. 36, 38, 39, 40, 41, 42). It thus appears critical to also compare the transgenerational effects of single versus multiple environmental changes, to test the predictability of complex parental effects and assess the meaningfulness of previous simplified studies. However, so far only few studies (e.g. 17, 23, 24, 43, 44) tested for the parental effects of multiple simultaneous environmental changes.

Here we used the model species *Arabidopsis thaliana* to thoroughly assess the generality and predictability of parental effects. We subjected multiple genotypes of *A. thaliana* to a broad range of biotic or abiotic environmental stresses, or combinations of these, altogether 24 different stress treatments, and then assessed phenotypic variation in the offspring of these plants. Our experimental set-up allowed us to address the following questions: (1) How common and how consistent are parental effects across different environmental stresses and plant genotypes? (2) Can the direction and magnitude of parental effects be predicted from the direct effects of environmental stresses on the parental generation? (3) Are the parental effects of multiple simultaneous environmental stresses additive or non-additive?

## Results and Discussion

### Generality and consistency of parental effects

Many of the studied abiotic or biotic environmental stresses, or their combinations, caused significant parental effects in our experiment. The magnitude and direction of these effects strongly depended on the treatment, plant genotype, and the measured plant trait (Table 1). The strongest parental effects were on plant biomass and fruit production, where several stresses experienced by mother plants increased or decreased the performance of their offspring by 30-40% (Figure 1). For instance, exposure of mother plants to cold, mild heat or shading transgenerationally increased biomass and reproduction by 20-35%, whereas intense heat, or salt in combination with drought, had the opposite effect and decreased both biomass and fruit production by similar amounts (Figure 1). The magnitudes of these effect sizes are well within the range of what previous studies have reported for parental effects in *A. thaliana* and other species (e.g. 6, 15, 16, 18, 26, 30, 45). Overall, 7 out of the 12 studied stresses had significant transgenerational effects on plant biomass, and 5 out of 12 on plant reproduction (Table 1). Thus, parental effects appear to be common in *A. thaliana*, and elicited by a broad range of environmental stresses – with likely consequences for ecological interactions and evolutionary trajectories (9, 10, 46).

**Figure 1.**
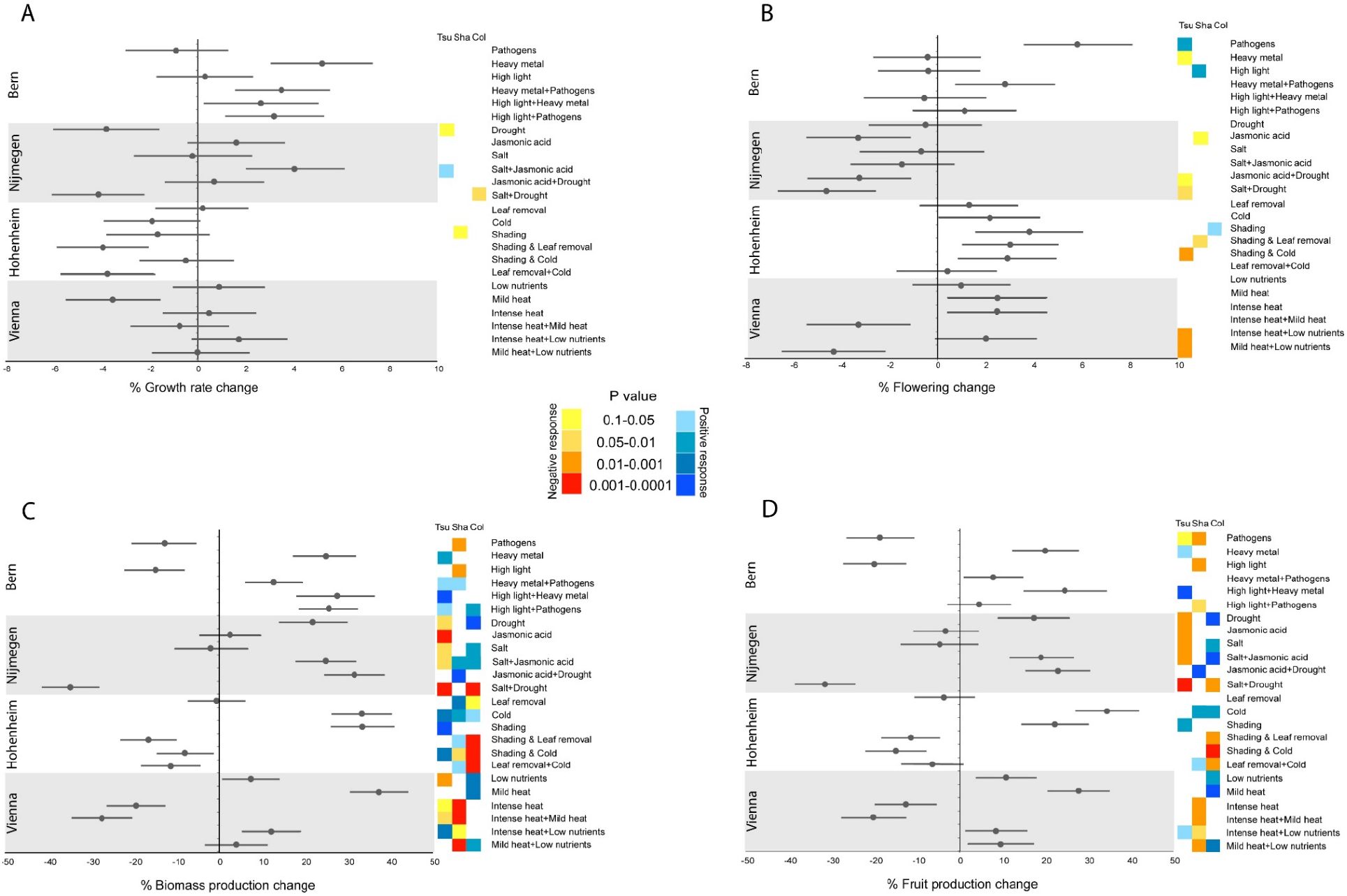
Effect sizes of parental effects of different environmental stresses, or their combinations, on *Arabidopsis thaliana* plants. The values are % differences (mean ± SE) in performance between the offspring of treated parents and the offspring of control parents. Note that the parental generation was grown in four different experimental locations. The coloured squares indicate the significance levels (from contrasts) of parental effects for individual genotypes (red spectrum = negative effects; blue spectrum = positive effects).

**Table 1.**
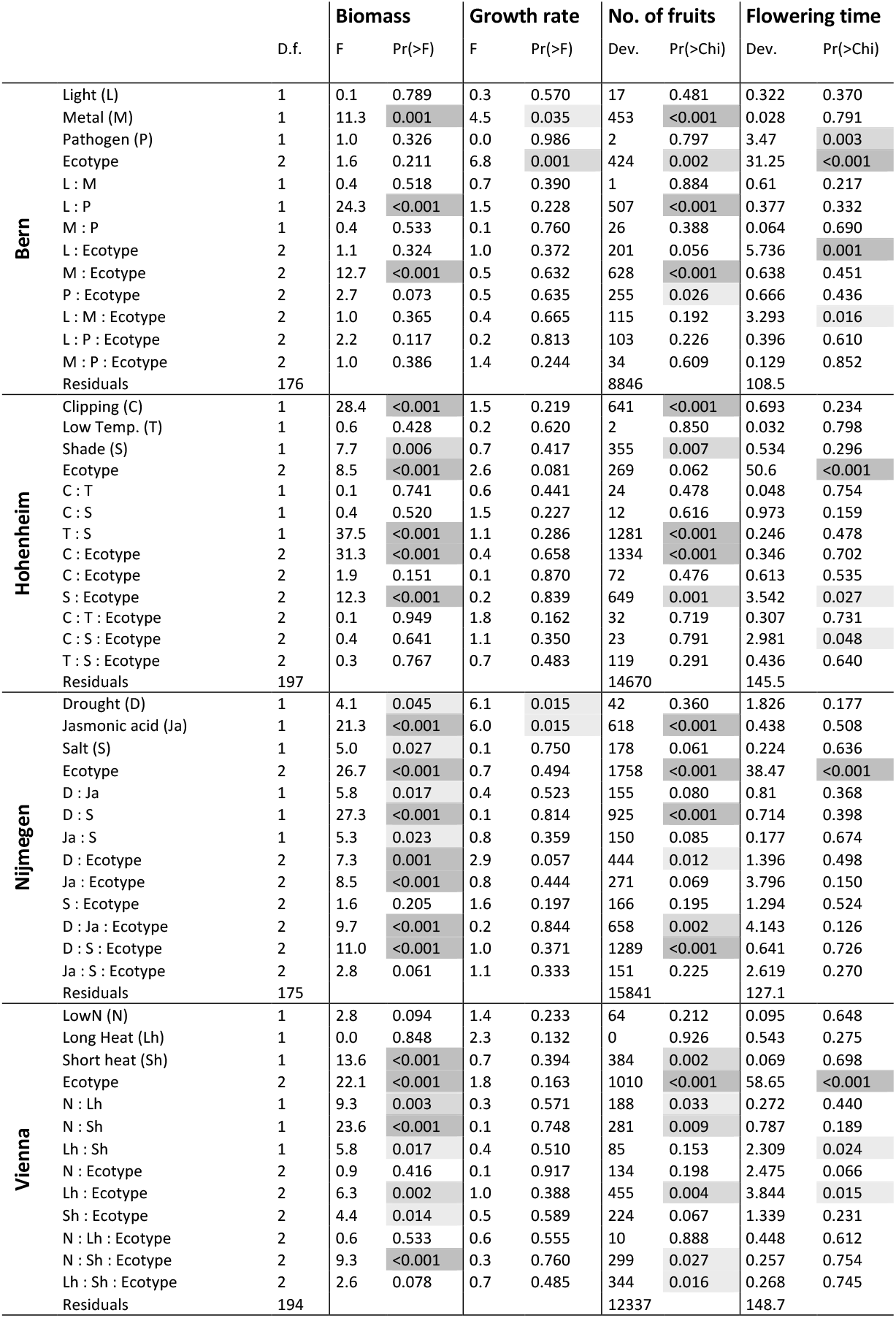
Results of ANOVA testing for parental effects of individual stresses, or their combinations, on the growth and fitness of three genotypes of *Arabidopsis thaliana*. Since the parental generation was grown in four different locations, the data were analysed separately for each. Effects significant at *P*<0.05 are highlighted.

Compared to plant biomass and reproduction, the growth rate and flowering time of plants were much less subject to parental effects, with only few percent changes across generations (Figure 1), and few individual stresses with significant transgenerational effects (Table 1). Clearly, some plant traits are much less prone to parental effects than others, possibly because they are under tighter developmental control. A good example is flowering time, which is strongly differentiated among geographic origins (significant ecotype effects in Table 1; see also 47), but it is hardly plastic across generations.

The three *Arabidopsis* ecotypes included in our study often differed in the degree and magnitude of transgenerational effects (Figure 1; significant ecotype interactions in Table 1). Sometimes the effects were even in opposite directions, resulting in non- or hardly significant main effects of an environmental stress across ecotypes. For instance, drought and salt stress had negative transgenerational effects (i.e. lower performance of offspring compared to the offspring of control plants) on the *Col* ecotype, but positive effects on *Tsu*, and none at all on *Sha* (Figure 1). Our results thus demonstrate substantial genetic variation for parental effects among *Arabidopsis* ecotypes, which supports previous studies with *Arabidopsis* and other plant species (e.g. 17, 29, 30, 48, 49, 50, 51; 52) that also found genotype-specificity of parental effects. Compared to previous studies, our experiment included a much broader range of environmental stresses, and it thus demonstrates that G x E effects are very common across generations, just as they are for within-generation plasticity (53, 54).

In summary, we find that parental effects are common and strong, but genotype-specific, in *Arabidopsis thaliana*. Because of this genotype-specificity, and their effects particularly on fitness-related traits, we should expect parental effects to influence selection and evolution of the species.

### Effects on parental versus offspring generation

Having demonstrated parental effects of a broad range of environmental stresses, we next asked if the direction and magnitude of these cross-generation effects was related to the within-generation effects of the different stresses. Intuitively, we expected that negative transgenerational effects would be caused by environmental stresses that also have negative effects on the same trait in mother plants, and vice versa. We found that this was the case for some environmental stresses. For instance, the combination of short intense heat with continuing mild heat significantly decreased the biomass of both mother plants and their offspring (Figure 2). However, there were also cases where within- and across-generation effects were in opposite directions. For instance, high light intensity increased the growth of mother plants, but it decreased offspring biomass, and for mild heat it was vice versa (Figure 2). There were also cases where stress treatments affected mother plants but not the offspring, e.g. for salt addition or intense heat, which strongly decreased the biomass of parents but had no effects across generations (Figure 2). Most interestingly, we observed also cases where the direct, within-generation effects of stresses were almost zero, but there were significant transgenerational effects. Examples are cold and drought, which did not at all affect the mother plants in our experiment, but they both strongly increased offspring biomass (Figure 2). Environmental stresses with strong direct impacts but no parental effects have been reported previously (e.g. 17, 25), but we are not aware of any previous studies that have shown the opposite. Altogether, because of the diversity of within-versus across-generation responses, there was no relationship between the stress responses of mothers and offspring in our experiment (*R^2^*=0.038, *P* = 0.358). While a discussion of the biological mechanisms underlying these diverse results is beyond the scope of this paper, an important take-home message is that the direction and magnitude of parental effects cannot be predicted from the parental responses to an environmental stress, and that sometimes seemingly ineffective environmental changes may nevertheless cause strong parental effects.

**Figure 2.**
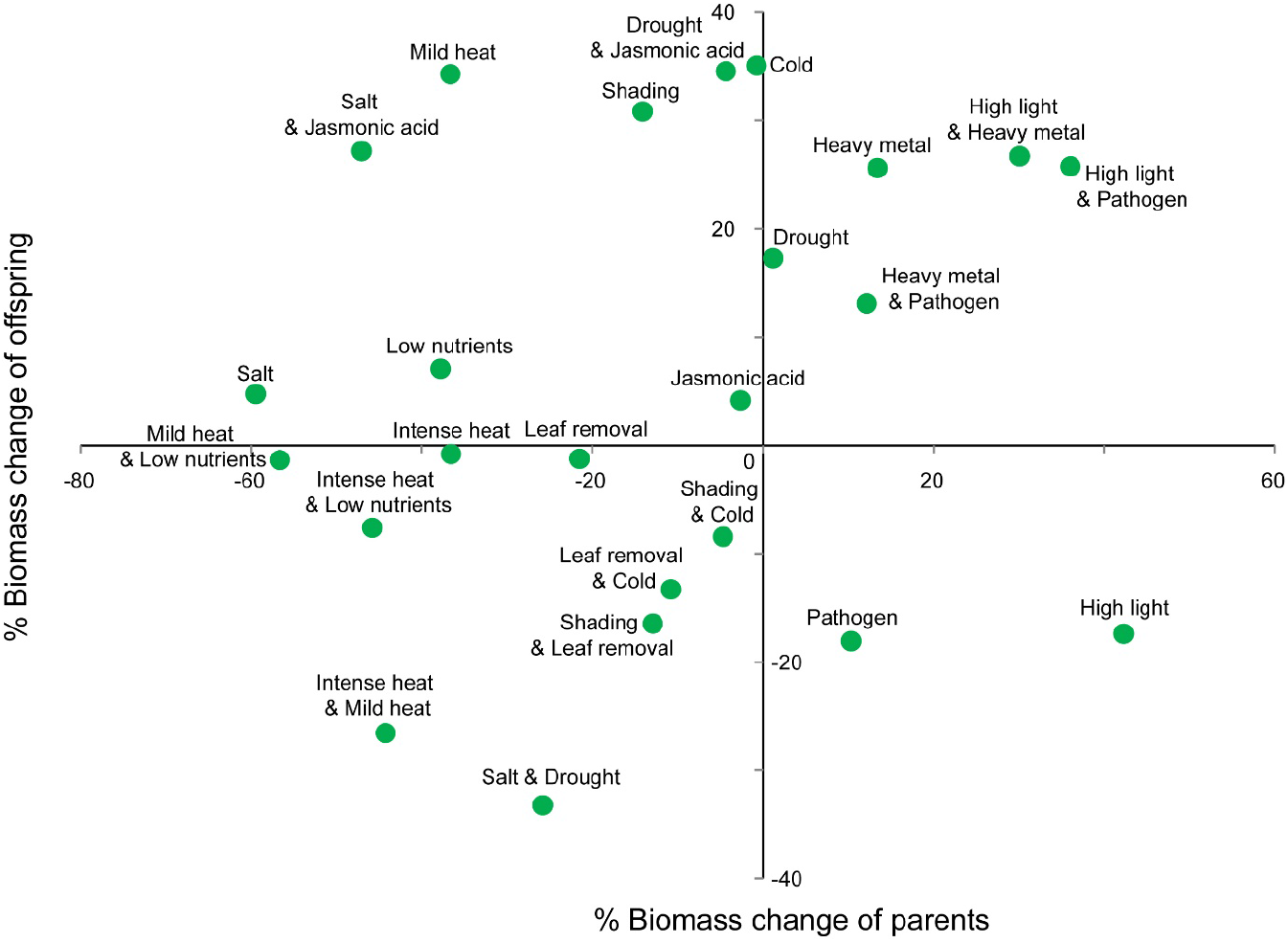
Relation of offspring biomass production responses to parental treatments with responses of parents to the treatments. The responses are % changes in biomass production of plants experiencing treatment (or offspring of parents of the treatments) in comparison to control plants (or offspring of control parents). Individual treatments are highlighted.

### Parental effects of multiple simultaneous environmental stresses

Environmental change is usually multifactorial (36, 37). It is therefore important to understand interactions between multiple drivers of environmental change, and their potential non-additive effects on organisms (e.g. 35, 38, 39, 41, 42). Our experiment allowed us to address these questions for parental effects of a broad range of environmental stresses on *A. thaliana*. We found that for 8 out of the 12 combinations of environmental stresses there were significant statistical interactions in their effects on plant biomass and/or fruit number (Table 1), indicating non-additivity of stresses when occurring in combination. For instance, high light intensity and pathogen infection caused negative parental effects on plant biomass when tested individually, but in combination they increased the biomass of offspring plants (Figure 1C). Positive parental effects of cold and shading turned into a negative effect when the two stresses were combined, and while drought and salt caused positive or neutral parental effects, their combination caused the strongest negative parental effect on plant biomass observed in our experiment (Figure 1C). In addition to the general interactions between environmental stresses, we also found several significant three-way interactions between two stresses and plant genotype (Table 1), i.e. the non-additivity of multiple stresses depended to some degree on the plant genotype. Our results corroborate the findings of the few previous studies that tested for transgenerational effects of multiple stresses (17, 24, 43) and that found similar non-additive effects. They clearly show that the non-additivity – or context-dependency – of multiple environmental stresses is another challenge for predicting parental effects, particularly under realistic conditions.

### Conclusions

In summary, our study demonstrates that parental effects strongly influence the growth and reproduction of *Arabidopsis thaliana*, and that many different environmental stresses can cause such parental effects. This is an important result also because we urgently need to understand the mechanisms by which plants respond to global environmental changes, and besides phenotypic plasticity (55, 56, 57) and longer-term adaptation (e.g. 58), parental effects might be another, somewhat intermediate, facet of plant responses. We also found that parental effects were strongly genotype-dependent, that effects of multiple stresses were often non-additive, and that there was no relationship between the within- and across-generation effects of environmental stresses. Thus, parental effects in *A. thaliana* are complex and difficult to predict, and we should be cautious with generalizing from simple studies with single plant genotypes and/or only few individual environmental stresses. From all we know about the ubiquity of G x E interactions, it seems likely that the situation is similar also for parental effects in other plant species. A thorough understanding of parental effects in plants will therefore be possible only with large experiments that include multiple plant genotypes and multiple, interacting environmental drivers.

## Methods

### Plant material

*Arabidopsis thaliana* is an annual species from open or disturbed habitats of the northern hemisphere. Because of its small genome size, predominant selfing and rapid life-cycle the species is a popular model species in plant biology as well as ecological and evolutionary genetics and genomics (59, 60). In our study we worked with three ecologically and geographically distinct genotypes of *A. thaliana*, the common laboratory strain *Col-0* (Versailles Center ID 168AV), the *Sha* genotype (VC ID 236AV) originating from Tajikistan and the *Tsu-0* genotype (VC ID 91AV) from Japan. All three genotypes are frequently used in genetics and plant biology, and have served as parents for populations of recombinant inbred lines. The same seed batch was used in all four experimental locations (see below).

### Parental generation

We subjected the plants to 12 different individual biotic and abiotic parental stress treatments, plus 12 pairwise combinations of these stresses, resulting in a total of 24 different stress treatments. For logistic reasons, the 24 treatments were distributed across four different labs (henceforth referred to as “locations”) in Bern, Hohenheim, Nijmegen and Vienna. In Bern, we tested the effects of light stress, heavy metal, pathogens, and all pairwise combinations of these. In Hohenheim, we tested the effects of cold treatment, shading and leaf removal (simulated grazing), and their combinations. In Nijmegen we tested the effects of drought, salt stress and jasmonic acid (simulation of herbivore attack), and their combinations, and in Vienna we tested two different kinds of heat stress, as well as the effects of low nutrients, and their pairwise combinations (see next section for more details on the treatments).

At each location, we grew the plants in temperature-controlled growth chambers under the same standardized temperature and daylength conditions (16/8h light/dark, 21°C/16°C), and we further minimized location differences by growing plants in the same pots (7 x 7 cm) and substrate (Einheitserde ED 63T) everywhere. We stratified seeds on wet filter paper at 4°C for three days and transplanted seedlings to individual pots. All plants were bottom-watered twice a week throughout the study. Sixteen days after sowing, we started the parental stress treatments, with six treatments (see above) plus a control treatment in each location, and seven replicates per treatment and genotype, i.e. 147 plants per location and 588 plants overall. Where possible, treatments were terminated when the plants started to bolt.

To estimate the growth rates of plants under different experimental conditions, we measured the rosette diameter of each plant at 16, 20, 24, 28 and 32 days after sowing, fitted a power function to each plant’s data, and used the parameter *b* as a measure of growth rate. Throughout the experiment, we continuously monitored plant phenology and recorded the date of first flowering (= first petals visible) of each plant. The plants were harvested sequentially, each at the same developmental stage when approximately one third of the siliques had reached maturity. We harvested each plant aboveground, counted its fruit number, and placed it in a paper bag for drying and after-ripening at room temperatures. After 14 days we collected the seeds from the paper bags, dried the remaining biomass at 70°C for 24 hours and weighed it. We pooled the seeds of all replicate plants per genotype and parental treatment and used these to establish the offspring generation (see below).

### Parental treatments

We experimentally subjected the parental plants to 12 different environmental treatments: (1) **Light stress** was imposed by increasing light levels from approximately 250 μmol m^-2^ s^-1^ in the control environment to 450 μmol m^-2^ s^-1^ in the treated plants. (2) **Heavy metal stress** was created by adding 5 ml of a 8 mMol solution of CuSO4 to each treated pot every second day, with the last addition at day 28 after sowing. (3) For **pathogen infection** we sprayed the plants four times (starting at day 16 after sowing, and then every third day) with a water solution containing 8 x 10^8^ bacteria of *Pseudomonas syringae* pv. *tomato* DC3000 per ml. The *P. syringae* DC3000 strain is strongly virulent and causes disease symptoms in *A.thaliana*. (4) **Cold stress** was imposed by regularly subjecting plants to 16 h of 4° C temperature during one week (16 h cold followed by 8 h at 21° C; a total of 112 h of cold). To keep plants at long-day conditions, the 16 h cold were divided into 8 h at light and 8 h at dark conditions. (5) **Shading** was created by growing plants under a shading filter foil (122 Fern Green; Lee Filters, Andover, UK) that reduced light by 50% and lowered the red:far red ratio to 0.2. The plants were kept shaded until the control plants began to flower. **(6) Leaf removal** was applied by cutting off all cotyledons, which at this time represented 50% of the leaf area, at day 16. 20 days later we repeated the treatment and again cut 50% of the leaf area of each plant. (7) **Drought stress** was created by not watering the treated plants unless they showed signs of wilting, whereas all other plants were watered regularly. (8) To create **salt stress,** we added a 4g/L NaCl solution at day 16 and after that treated plants twice a week with a 8 g/L NaCl solution until day 30. (9) **Jasmonic acid** was applied by spraying treated plants with a 0.5 mM jasmonic acid solution (Cipollini et al. 2002) and control plants with a mock treatment of 0.5% ethanol every second day starting at day 16 days after sowing. (10) **Low nutrient stress** was created by transplanting plants into a nutrient-poor substrate (Huminsubstrat N3, Neuhaus, Germany) instead of the standard substrate used for all other plants. (11) **Short intense heat** stress was created by moving plants for 24 h to a 37°C growth chamber at day 16 and then back to control conditions, whereas for the (12) **prolonged mild heat** treatment plants were moved to a 30°C growth chamber for 10 days, starting at day 16. For the combination of the two heat treatments, the plants were first moved to the 37° chamber for 24 h and then to the 30°C chamber for another nine days.

### Offspring phenotyping

To test for the effects of parental stress treatments, or their combinations, on offspring phenotypes, we used the seeds collected from the parental generation to grow offspring of all genotypes and parental treatments in a common greenhouse environment. Using the same protocols for germination and growth and the same pots and substrate as for the parental generation, we grew 10 replicate plants per genotype and treatment (= a total of 24 x 3 x 10 = 720 plants) in a greenhouse with a 16/8 h light/dark cycle and temperatures of 27/16°C (day/night). The plants were arranged in a fully randomized order and watered regularly. We measured the same phenotypic traits as in the parental generation: growth rate, aboveground biomass, flowering time, and fruit number.

### Statistical analyses

The parental generation data were analysed through linear models in which we tested the effects of stress treatments, plant genotype, and their interactions, on the growth rate, aboveground biomass, fruit production and flowering time of plants. We carried out these analyses separately for each of the four locations.

For the offspring generation data, we first examined how large the differences between the four parental locations were, in spite of our efforts to standardize conditions. A two-way ANOVA testing for location and genotype effects among the control plants only showed that there were still large differences among locations (*P* < 0.001 for all traits), and we therefore decided to also analyse the offspring data separately for each location. We used similar linear models as for the parental generation analyses, testing for the effects of parental treatment and genotype, and their interactions, in each location. If the main effect of parental treatments was significant, we additionally tested a series of contrasts comparing each treatment combination to the control group, to identify which specific parental treatments had significant effects on the offspring. We first ran these analyses across all genotypes and then, since genotype by treatment interactions were significant in most cases, also separately for each genotype.

To test for a relationship between the magnitude and direction of parental and offspring stress responses, we calculated the cross-genotype % change caused by each treatment when compared to the respective control plants. We did this for the parental and offspring data and then used linear regression to test for a relationship between the two.

## Acknowledgements

This work, as part of the European Science Foundation EUROCORES Programme EuroEEFG, was funded by the Swiss National Science Foundation (SNF grant 31EE30-131171 to O.B. and M.F.), the German Research Foundation (DFG grant SCHM 1354/4-1 to K.S., the Austrian Science Fund (FWF I489 to Ortrun Mittelsten Scheid, and FWF I3687 to R.G.), and the Dutch Research Council NWO grant to J.O.). V.L. acknowledges support from the Czech Science Foundation 20-00871S and RVO 67985939. We are extremely grateful to Vincent Colot and Ortrun Mittelsten Scheid for their intellectual contributions to this study.

